# FMN-dependent oligomerization of putative lactate oxidase from *Pediococcus acidilactici*

**DOI:** 10.1101/790964

**Authors:** Yashwanth Ashok, Mirko M. Maksimainen, Tuija Kallio, Pekka Kilpeläinen, Lari Lehtiö

## Abstract

Lactate oxidases belong to a group of FMN-dependent enzymes and they catalyze a conversion of lactate to pyruvate with a release of hydrogen peroxide. Hydrogen peroxide is also utilized as a read out in biosensors to quantitate lactate levels in biological samples. *Aerococcus viridans* lactate oxidase is the best characterized lactate oxidase and our knowledge of lactate oxidases relies largely to studies conducted with that particular enzyme. *Pediococcus acidilactici* lactate oxidase is also commercially available for e.g. lactate measurements, but this enzyme has not been characterized before in detail. Here we report structural characterization of the recombinant enzyme and its co-factor dependent oligomerization. The crystal structures revealed two distinct conformations in the loop closing the active site, consistent with previous biochemical studies implicating the role of loop in catalysis. Despite the structural conservation of active site residues when compared to *Aerococcus viridans* lactate oxidase we were not able to detect either oxidase or monooxygenase activity when L-lactate or other potential alpha hydroxyl acids were used as a substrate. *Pediococcus acidilactici* lactate oxidase is therefore an example of a misannotation of an FMN-dependent enzyme, which catalyzes likely a so far unknown oxidation reaction.

## Introduction

Alpha-hydroxy acids are oxidized by a family of FMN-dependent enzymes [1]. Lactate oxidases belong this class of enzymes along with other well characterized members such as lactate monooxygenase, glycolate oxidase, flavocytochrome b2 (L-lactate dehydrogenase). Lactate oxidases catalyze the conversion of lactate to pyruvate and H_2_O_2_. *Aerococcus viridans* lactate oxidase (AvLCTO) is a prototypical example of lactate oxidase. AvLCTO is a biotechnologically important enzyme used in manufacturing biosensors [2,3], that measure L-lactate levels in biological samples using an indirect assay that utilizes H_2_O_2_ released as a by-product.

Catalysis of L-lactate to pyruvate by lactate oxidases is thought to occur in two half reactions: (1) Binding of L-lactate results in reduction of FMN. Based on experimental evidence, His265 deprotonates the substrate from α-carbon, eventually leading to hydride transfer from substrate to N5 of FMN. This is the reductive half of the reaction. (2) Molecular oxygen oxidizes FMN to continue the catalytic cycle, which forms the oxidative half of the reaction. Lactate monooxygenases differ from lactate oxidases only in their oxidative half reaction. Lactate monooxygenases undergo ‘coupled’ reaction wherein the α-keto acid has a longer residence time in the active site, allowing reaction with hydrogen peroxide to undergo oxidative decarboxylation of pyruvate to produce acetate, carbon dioxide and water. This contrasts with lactate oxidases which undergoes ‘uncoupled’ pathway, where pyruvate is dissociated immediately before hydrogen peroxide reacts with pyruvate [4,5].

Both AvLCTO and *Pediococcus acidilactici* lactate oxidase (PaLCTO) are commercially available enzymes purified from native sources that are used for biosensor applications [6]. While AvLCTO has been characterized extensively using recombinant system, PaLCTO has not been studied. PaLCTO is annotated as lactate oxidase and the closest homolog in *Pediococcus acidilactici* proteome does not use lactate as a substrate. We describe here structural and biophysical studies of PaLCTO, which revealed FMN-dependent folding and oligomerization of the enzyme.

## Materials and methods

### Cloning and site directed mutagenesis

*Pediococcus acidilactici* proteome wide search for keywords lactate, lactate oxidase, α-hydroxy acid in Uniprot database gave only one hit that matched with AvLCTO. Similarly, BLASTp search with AvLCTO against *P. acidilactici* DSM20284 revealed the same hit annotated as putative L-lactate oxidase (Uniprot E0NE46). Genomic DNA of *Pediococcus acidilactici* (DSM 20284) and *Aerococcus viridans* (DSM 20340) was obtained from DSMZ, Germany. Gene encoding PaLCTO was cloned into pNH-TrxT (Structural Genomics Consortium) vector using SLIC cloning. The vector encodes for an N-terminal thioredoxin tag with a cleavable TEV protease recognition site. A94G mutant was obtained using standard site-directed mutagenesis protocol. All clones were verified using Sanger’s dideoxy sequencing. Untagged PaLCTO and AvLCTO were recombinantly expressed from pNIC-CH vector with a stop codon added immediately after the native sequence. Commercial PaLCTO was purchased from Sigma (catalog number LO638, lots STBG2905V and STBF3223V).

### Protein expression and purification

Proteins were expressed using BL21 (DE3). Overnight culture was inoculated into terrific broth auto-induction media supplemented with 8 g/L glycerol and 50 μg/ml kanamycin. Cells were grown at 37°C until OD_600_ reached 1. The temperature was reduced to 18°C for overnight for protein expression. Cells were harvested the next day by centrifugation and either directly used for purification or flash frozen.

Cells were resuspended in lysis buffer (50 mM Hepes pH 7.5, 0.5 M NaCl, 10% glycerol, 0.5 mM TCEP, 10 mM imidazole) with 0.1 mM Pefabloc SC (Sigma-Aldrich) and lysed at 20 kPsi using cell disruptor (Constant systems, UK). Lysates were cleared by centrifugation at 16,000 rpm for 30 minutes. Supernatant was loaded on 4 ml of HisPur™ Ni-NTA resin (Thermo Scientific). The beads were washed with 10 ml of lysis buffer and then with 15 ml of wash buffer (50 mM Hepes pH 7.5, 0.5 M NaCl, 10% glycerol, 0.5 mM TCEP, 25 mM imidazole) was used to capture recombinant proteins. Proteins were eluted with elution buffer (50 mM Hepes pH 7.5, 0.5 M NaCl, 10% glycerol, 0.5 mM TCEP, 350 mM imidazole). Buffer exchange was achieved using Amicon 30 kDa concentrator and the tags were cleaved by digestion with TEV protease (in-house preparation) for 2 days at 4°C (7). Supernatant was passed again through Ni-NTA resin to capture thioredoxin and TEV protease. Tag cleaved PaLCTO was collected from flow through. The proteins were purified subsequently with HiLoad 16/600 Superdex 200 using size exclusion buffer (30 mM Hepes pH 7.5, 0.35 M NaCl, 10% glycerol, 0.5 mM TCEP). Proteins were concentrated and were flash frozen in liquid N_2_ and stored at - 70°C. Purified PaLCTO showed bright yellow color due to the presence of a co-factor.

For purification of untagged PaLCTO and AvLCTO, cells were resuspended in 20 mM Potassium Phosphate pH 7.0 with 0.1 mM Pefabloc and lysed as described above. Clarified lysate was loaded into Q sepharose, followed by washing with lysis buffer. Proteins were eluted using 0-1 M KCl gradient in lysis buffer. Fractions containing required proteins were pooled and solid ammonium sulfate was added to a concentration of 1.5 M. After 15-minute incubation on ice precipitates were discarded using centrifugation. Proteins were loaded to phenyl sepharose and washed with 20 mM Potassium phosphate, 1.5 M ammonium sulfate and were eluted using a gradient from 1.5 M ammonium sulfate to 0 M in 20 mM Potassium Phosphate pH 7.0. Pooled from phenyl sepharose were loaded to Superdex200 16/600 equilibrated with 100 mM Potassium phosphate pH 7.0.

For FMN incorporation experiments, 1:2 molar ratio (protein:FMN) was used and incubated on ice for overnight and subsequently purified using SEC in Superdex200 16/600 using 30 mM Hepes pH 7.5, 0.35 M NaCl, 10% glycerol, 0.5 mM TCEP.

### Analytical size exclusion chromatography (aSEC)

Bio-Rad gel filtration standards were used for calibrating Superdex 200 Increase 10/300 column. 30 mM Hepes pH 7.5, 350 mM NaCl, 0.5 mM TCEP was used as buffer with 0.5 ml/min flow rate. Blue dextran was used for void volume calculation. Protein elution volumes were calculated using peak elution time with absorption at 280 nm. Partition coefficient (K_av_) were calculated using elution volumes with the equation K_av_ = (V_e_-V_0_)/ (V_t_-V_0_), where V_e_ is the elution volume, V_0_ is the void volume and V_t_ is the total volume of column.

### Deflavination

Protein aliquots were thawed and diluted in 250 mM sodium phosphate pH 7.5, 3 M Potassium bromide, 10% glycerol, 0.5 mM TCEP, 1 mM EDTA and left on ice for 4 days [8]. Following incubation, solution was concentrated using a 10 kDa cut-off concentrator. Majority of the yellow color was in the flow through. Deflavination was repeated to completely remove FMN and the yellow color. Pure monomeric proteins were obtained using size exclusion chromatography using size exclusion buffer.

### CD spectroscopy

CD spectra were recorded using Chirascan™ spectrophotometer (Applied photophysics Ltd, USA). Spectra scan experiments were conducted at 22°C using a 0.1 cm optical path length cuvette. Proteins were diluted to 0.1 mg/ml in water for measurements. Spectra were recorded from 190-280 nm in 1 nm steps thrice and averaged for each sample. Thermal stability measurements were done by heating the protein 22-80°C. Data analysis was carried out by using pro-data viewer (Applied Photophysics Ltd, USA). Data obtained at 222 nm was used for calculating melting temperature using Boltzmann sigmoidal equation in Graphpad Prism.

### Crystallization

Crystals were readily obtained in several conditions from the initial crystallization trials using the JCSG+ and PACT premier™ crystallization screens (Molecular Dimensions). Crystals used for data collection were obtained in 24 hours either in a hanging or sitting drops, which were prepared by mixing 75-150 nl of 15 mg/ml protein with 75-150 nl of 1.1 M Sodium Malonate, 0.1 M Hepes pH 7.0, 0.5% Jeffamine ED-2003 at 20°C.

### Data collection, processing & refinement

The crystals were cryoprotected with a solution containing 20% (v/v) glycerol in the mother liquor and flash-frozen in liquid nitrogen. All measurements were carried out at 100 K. WT PaLCTO dataset was collected at ID30A-1 and refolded dataset was collected at ID30B (European Synchrotron Radiation, France). Mutant dataset was collected at I04 (Diamond Light Source, UK). Diffraction data were indexed, scaled and merged using XDS [9]. Phases for the wild type PaLCTO were solved using molecular replacement with Phaser MR of the Phenix package [10,11] using AvLCTO (2J6X) as a search model [12]. Iterative model refinement was carried out using Refmac5 in CCP4 [13]. Model building and visualization were performed using Coot [14,15]. Structure related figures were generated using Pymol (Schrödinger). The data collection and structure refinement statistics are presented in Table 1.

**Table 1.** Data collection and refinement statistics for the crystal structures.

| Structure | Native WT PaLCTO | A94G PaLCTO | Refolded WT PaLCTO |
| --- | --- | --- | --- |
| PDB code | 6RHT | 6RHV | 6RHS |
| Beamline | MASSIF 1, ESRF | I04, DLS | ID30B, ESRF |
| Wavelength (Å) | 0.980 | 0.97 | 0.96770 |
| Space group | I422 | I422 | I422 |
| Cell dimensions a, b, c (Å) | 135.24 135.24 124.84 | 135.53 135.53 124.86 | 135.50 135.50 125.69 |
| Resolution (Å) | 95.63 (1.9) | 50 (2.0) | 50 (2.6) |
| R<sub>merge</sub> | 16.0 (114) | 19.5 (123.3) | 21.9 (131.6) |
| I / σI | 10.3 (2.2) | 11.98 (2.17) | 11.70 (2.19) |
| Completeness (%) | 99.9 (100) | 100 (100) | 100 (100) |
| Redundancy | 8.9 (9.1) | 13.2 (13.6) | 13.3 (13.3) |
| **Refinement** |  |  |  |
| R<sub>work</sub> / R<sub>free</sub> | 0.182 / 0.211 | 0.148 / 0.173 | 0.186/ 0.216 |
| **No. atoms** |  |  |  |
| Protein | 2895 | 2800 | 2743 |
| Ligand FMN | 31 | 31 | 31 |
| Water | 124 | 236 | 20 |
| **B-factors** |  |  |  |
| Protein | 29.62 | 25.25 | 40.3 |
| Ligand FMN | 23.2 | 21.74 | 31.3 |
| Glycerol | 52.05 | 48.64 | 54.77 |
| Water | 30.96 | 34.95 | 32.34 |
| **R.m.s.d.** |  |  |  |
| Bond lengths (Å) | 0.010 | 0.009 | 0.009 |
| Bond angles (°) | 1.628 | 1.54 | 1.675 |
| **Ramachandran plot (%)** |  |  |  |
| Favored regions | 96.69 | 97.28 | 96.34 |
| Additionally allowed regions | 3.04 | 2.45 | 3.38 |
| Outliers | 0.28 | 0.27 | 0.28 |

### Small angle X-ray scattering

SAXS was conducted in size-exclusion chromatography mode. Scattering contributions due to buffer were subtracted from protein peak using ScÅtter. SAXS based molecular weight estimates were done using SAXS MoW 2.0 [16]. Analysis of crystal structure fitting to scattering profiles was done using CRYSOL [17,18].

### Activity assays

Oxidase activity towards L-lactate and glycolate was measured in reaction mixture containing aminoantipyrine (4-AAP) and N,N-dimethylaniline (DMA) that react to form quinonediimine dye when they are oxidized by the horseradish peroxidase enzyme (HRP) and hydrogen peroxide generated in lactate oxidase reaction [4]. The initial reaction cocktail contained for each reaction 40 µl of 200 mM 3,3 dimethylglutaric acid-NaOH buffer, pH 6.5 (DMGA), 20 µl HRP solution (50 U/ml, Sigma-Aldrich P-8250), 20 µl 4-AAP and 30 µl deionized ELGA water. The cocktail was mixed by inversion and 110 µl was pipetted into microplate wells. Next, 25 µl of substrate solution and 40 µl DMA 0.2 % (v/v) solution were added into the wells. The microwell plate was transferred to a reader (Varioskan Flash or Tecan infinite pro) and incubated at +37 °C for 5 min. Finally, 25 µl of enzyme dilution (∼ 2 µg/ml) in 10 mM sodium phosphate buffer containing 0.1 mM flavin mononucleotide co-factor (FMN, Sigma-Aldrich F2253) was dispensed into the wells and absorbance at 565nm was monitored with one minute intervals for 15 minutes. Oxidase activity was calculated by dividing the slope of the linear regression line of enzyme activity graph (A565nm) from t = 0 to t = 15 min by the amount of enzyme protein taken to the assay. Standard L-lactate concentration in the assay was 1 mM, but when enzyme activities were screened, concentrations used ranged from 1 mM to 62.5 mM, and protein amount in the assay was increase even 250-fold. In K_m_ determinations, L-lactate concentrations were from 0.2 mM to 2.0 mM. Instead of FMN, flavin adenine dinucleotide (FAD) was tested as a co-factor in some enzyme assays.

For lactate 2-monooxygenase assay, protein samples were incubated in a similar mixture as previously, but 4-AAP and DMA were omitted. In addition to Na-phosphate buffer, 10 mM imidazole-HCl, 150 mM NaCl (pH 7.0) and 30 mM Hepes, 150 NaCl (pH 7.0) buffers were tested, since phosphate inhibits some enzymes using lactate as a substrate. All three buffers contained also 10 mM FMN. The assays were incubated overnight at +37 °C. Active lactate oxidases were used as controls to demonstrate detection and quantitation of activity. After incubation, reaction mixtures were analyzed by capillary electrophoresis. The mixtures were filtered with a 0.45 µm GHP Acrodisc syringe filter prior to analysis that were carried out with a P/ACE MDQ CE instrument (Beckman-Coulter, Fullerton, CA, USA) equipped with a diode array detector (DAD) using uncoated fused-silica capillaries of I.D. 25 µm and length 30/40 cm (effective length/total length). The samples were injected at a pressure of 3447.4 Pa for 10 s with a separation voltage of +16000 V. Calibration curves were created for external quantification. In addition, sample runs were performed also with spiked standards to confirm the identity of the analytes. Similar capillary electrophoresis assay was used also to test oxidase activity of PaLCTO towards 4-hydroxy mandelate.

## Results

### Characterization of purified protein

Recombinant enzyme was produced in *E. coli* and purified using Ni^2+^-NTA affinity chromatography. During subsequent size exclusion chromatography, PaLCTO eluted as two peaks. The first peak was yellow in color and the second was colorless (S1 Fig). Mass spectrometric analysis of the proteins present in the fractions revealed that both of the peaks actually contained PaLCTO. Previous reports demonstrate that the other members of this enzyme family are oligomeric and in order to study the oligomerization properties of the protein in both fractions, analytical size exclusion chromatography was performed with molecular weight standards (Fig 1A). The first peak containing the yellow fraction was judged to be tetrameric with a molecular weight of 162 kDa, while the colorless protein was monomeric with a molecular weight of 33 kDa. These results are in good agreement with the theoretical molecular weight of a monomer (39 kDa) and a tetramer (156 kDa).

**Fig 1.**
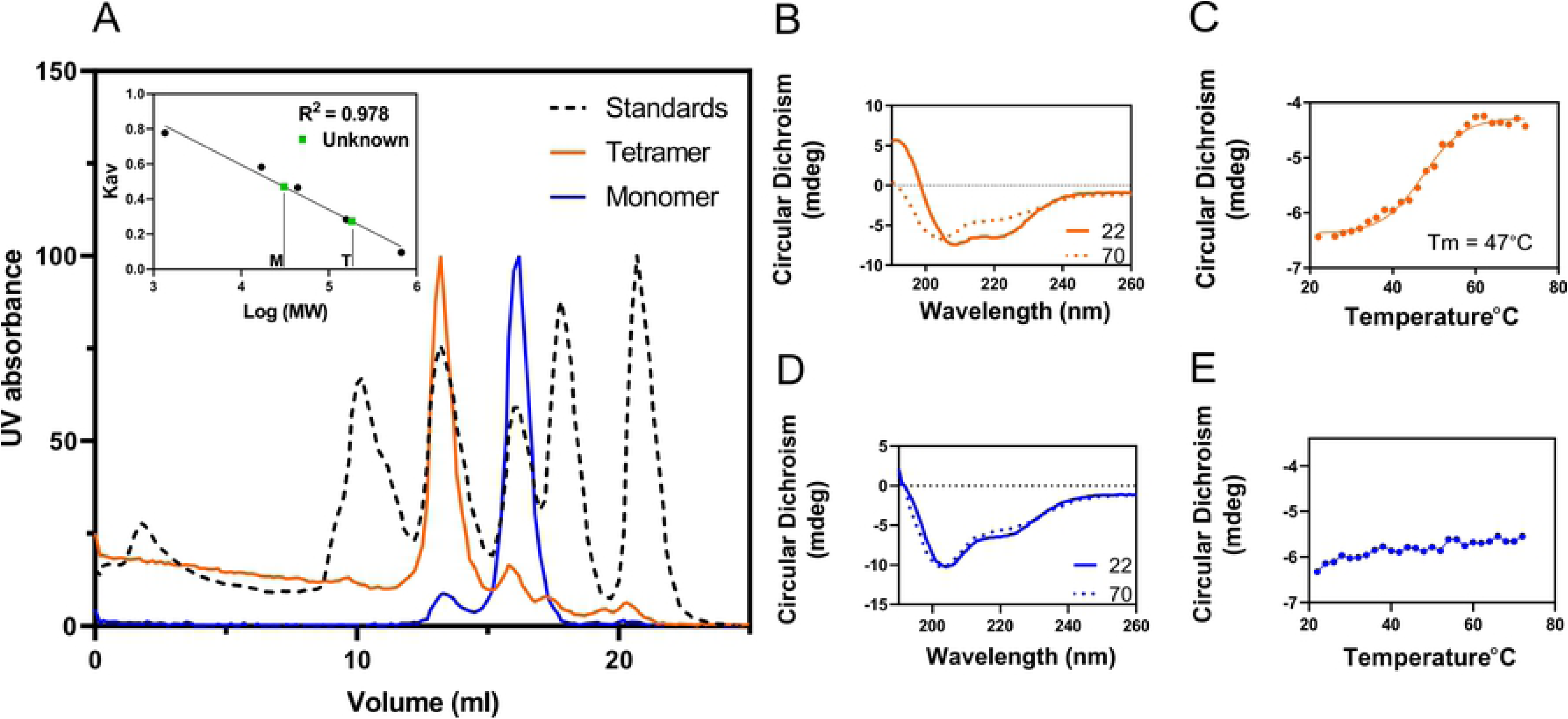
Characterization of purified PaLCTO. A) analytical size exclusion chromatography of purified tetrameric and monomeric fractions of PaLCTO. (inset: calibration curve. M-Monomer T-tetramer). B) CD spectra of tetrameric PaLCTO at 22 and 70°C. C) Melting temperature analysis of tetrameric PaLCTO at 222 nm. D) CD spectra of monomeric PaLCTO at 22 and 70°C. E) Melting temperature analysis of monomeric PaLCTO at 222 nm.

Flavin mononucleotide (FMN) is the co-factor that imparts bright yellow color to the protein and since this is absent in the monomeric protein, circular dichroism (CD) spectroscopy was performed to assess any possible differences in secondary structure between tetrameric and monomeric proteins. Tetrameric fraction showed typical features of a folded protein rich in secondary structures (Fig 1B). Negative peak maxima at 202 and 222 nm and a positive peak at 195 nm indicated that the protein has an alpha-helical content. Monomeric protein showed negative peak maxima at 204 nm, indicating that the monomeric protein showed no typical characteristics of a folded protein (Fig 1D). Melting curve studies using CD were performed to determine if the proteins are folded. Tetrameric fraction clearly showed sigmoidal transition at 222 nm whereas monomeric fraction did not show any such transition (Figs 1C and 1E). The calculated melting temperature of the tetrameric protein was 47°C. These results indicated that the monomeric protein has little or no secondary structural elements typical for a folded protein.

We also performed small angle X-ray scattering (SAXS) analysis for both of the proteins to further strengthen data obtained from CD and aSEC (Figs 2A and 2D). Analysis of raw scattering curves from tetrameric and monomeric fractions with SAXS MOW 2.0 indicated that the proteins had a molecular weight of 153 and 40 kDa, respectively. Normalized Kratky plot analysis revealed that both of the proteins were actually globular (Figs 2B and 2E), which was surprising in the light of the CD analysis of the monomer (Fig 1D). For a completely disordered protein one would expect the Kratky plot to show a constant rising in increased angle and therefore it appears that the monomer stays in a globular form despite that it does not contain distinct secondary structures. Distance distribution analysis shows Dmax values for the tetrameric and monomeric protein to be 113 Å and 66 Å, respectively (Figs 2C and 2F) and the tetramer Dmax correlates well with the crystal structure of AvLCTO. A summary of SAXS parameters obtained is summarized in (S1 Table and S2 Fig).

**Fig 2.**
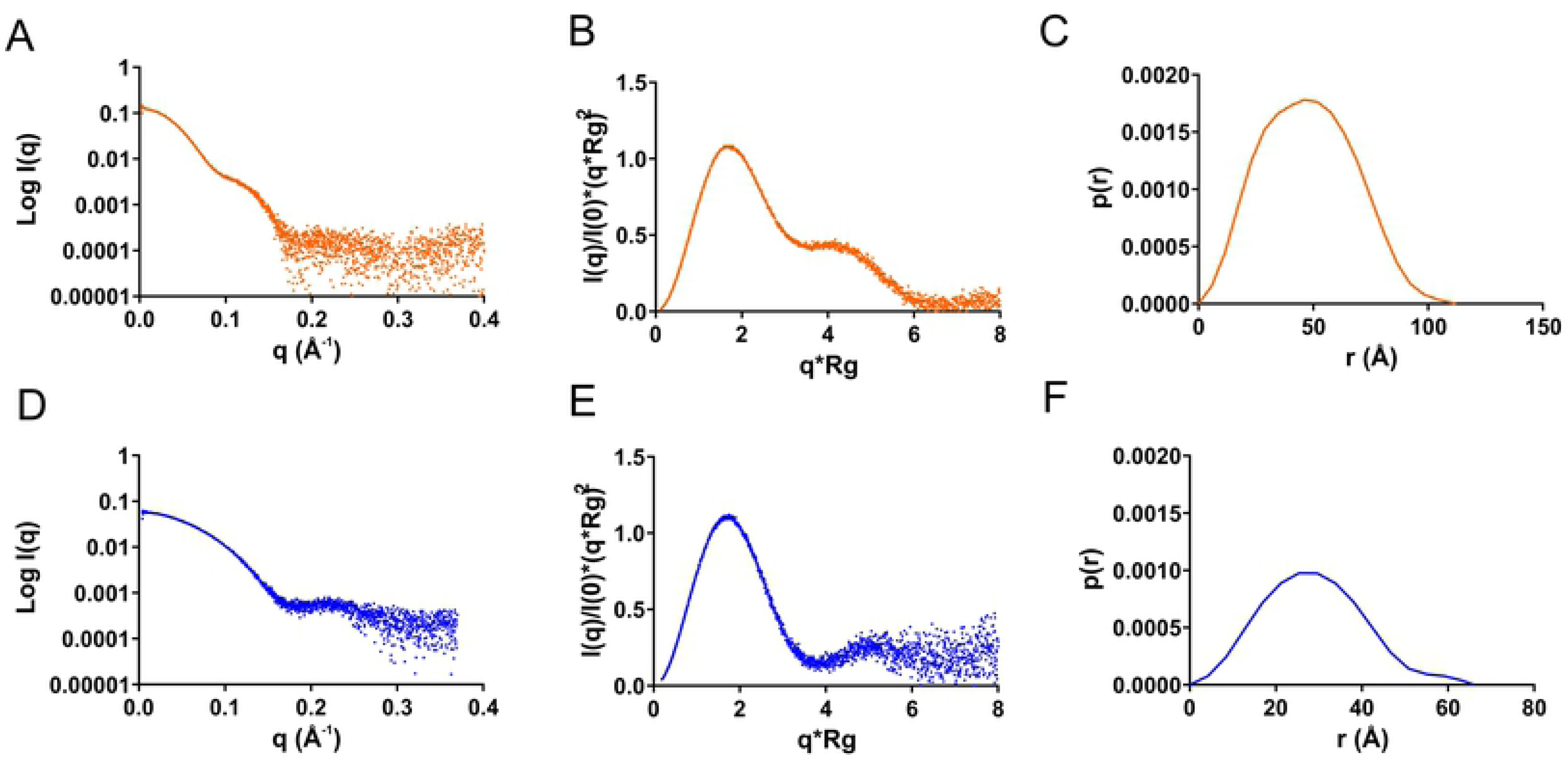
SAXS analysis of tetrameric and monomeric PaLCTO. A) Experimental scattering profile. B) Normalized Kratky plot and C) P(r) distribution curves for tetrameric native PaLCTO. D) Experimental scattering profile. E) Normalized Kratky plot and f) P(r) distribution curves for monomeric PaLCTO.

### Co-factor dependent oligomerization

In order to study whether monomeric enzyme could incorporate the co-factor, we incubated it with FMN and subsequently carried out SEC analysis to see if the protein oligomerizes in the presence of the co-factor. The protein indeed eluted at the same elution volume as native tetrameric protein (Fig 3A) indicating co-factor dependent oligomerization. Previously this phenomena has been described for some FAD containing enzymes [19,8,20]. We also performed loss of FMN binding studies, where native tetrameric proteins were chemically treated to remove FMN and the deflavination resulted in monomeric proteins (Fig 3B).

**Fig 3.**
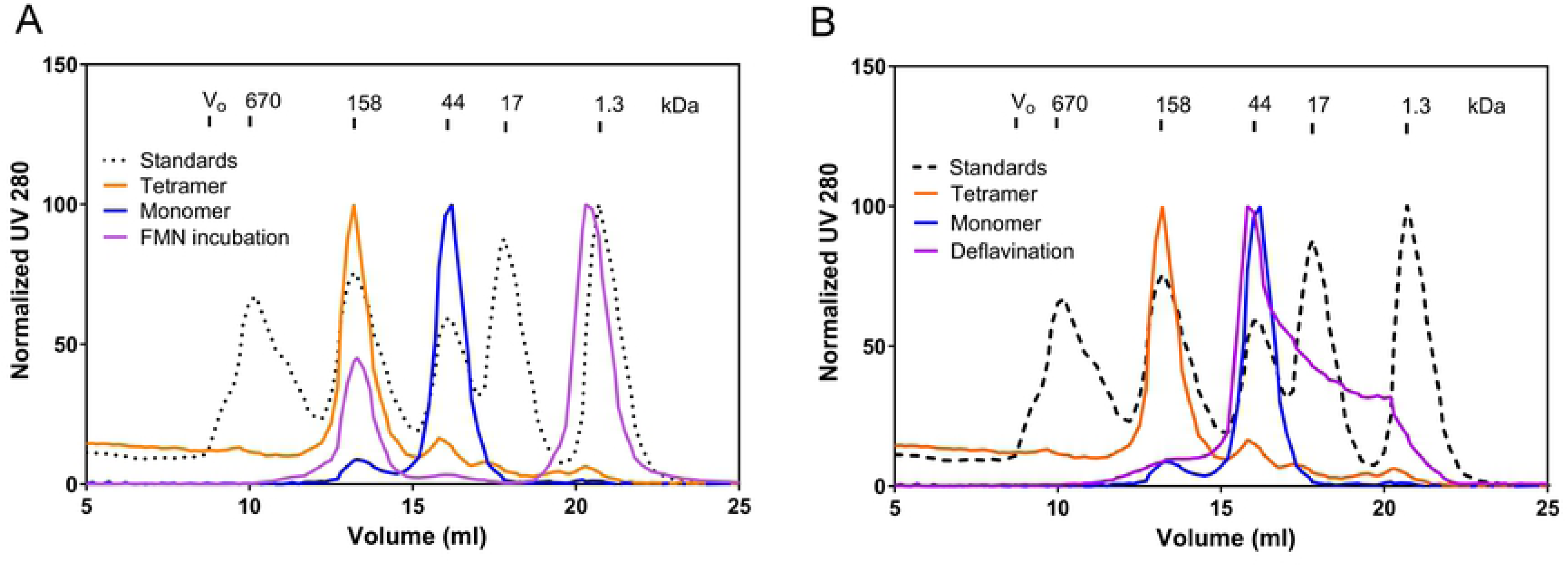
FMN dependent oligomerization of PaLCTO. A) Monomeric PaLCTO upon incubation with FMN forms a tetramer that elutes at the same elution volume as native PaLCTO. B) Deflavination of native PaLCTO makes it monomeric causing it to elute in monomer elution volume.

To assess if refolded proteins have similar secondary structure to native PaLCTO, we performed CD spectroscopic studies. Refolded protein had similar spectra to WT native PaLCTO and melting temperature (47°C) (S3A Fig and Fig 4A). Together the results show that PaLCTO could be reversibly folded and unfolded *in vitro*. SAXS analysis of refolded PaLCTO (Figs 4B and 4C) were also in line and indicated that the protein had a molecular weight of 151 kDa, Kratky plot was similar to native PaLCTO (Fig 4C) and Dmax was 113 Å as for the native tetramer (Fig 4D and S1 Table).

**Fig 4.**
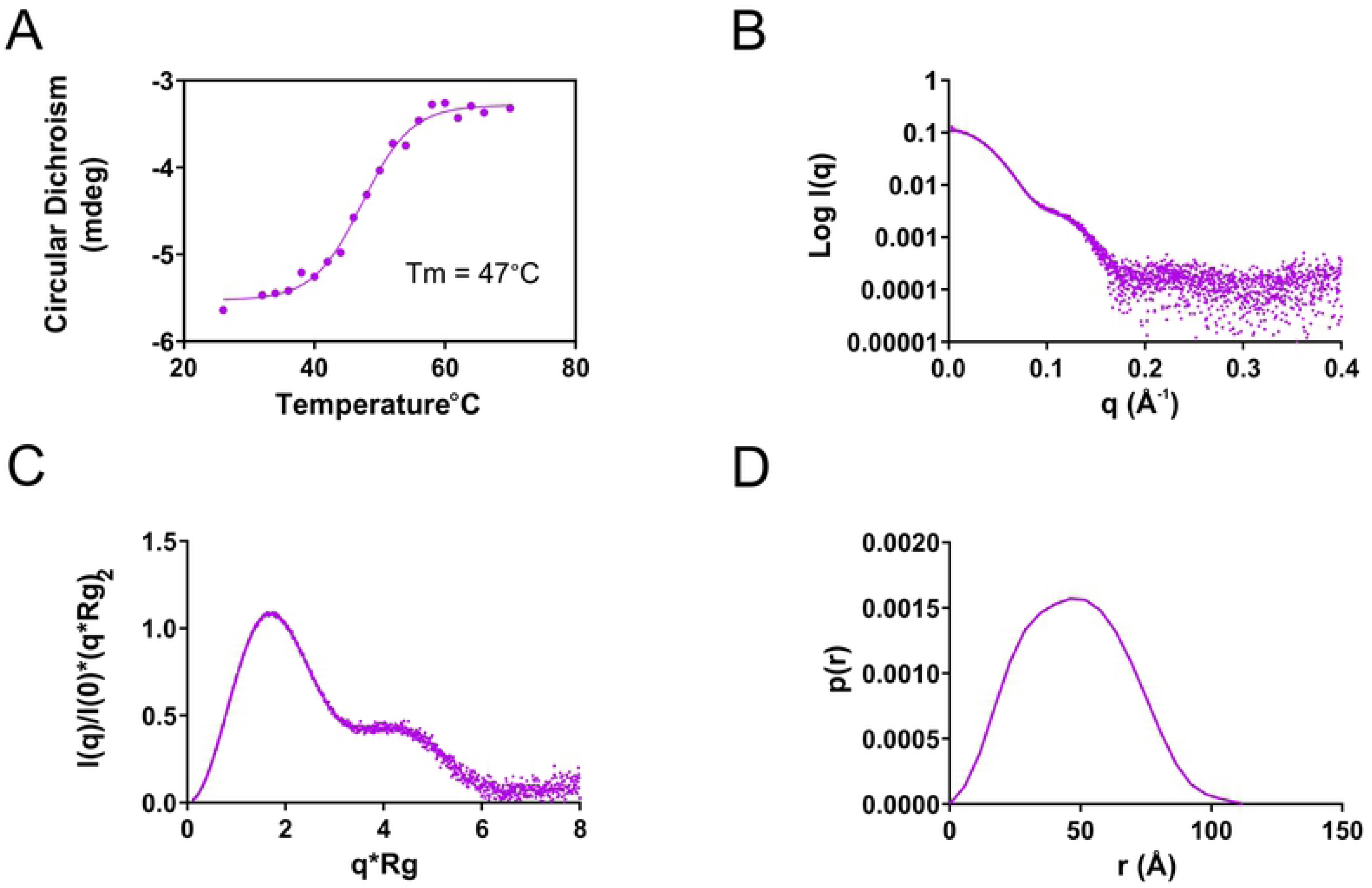
Analysis of refolded PaLCTO. A) Melting temperature analysis of refolded PaLCTO using CD from 222 nm. B) Experimental scattering profile. C) Normalized Kratky plot of refolded enzyme. D) P(r) distribution of refolded PaLCTO.

For AvLCTO, although it is well characterized, cofactor-dependent oligomerization has not been reported. Recombinant AvLCTO showed only one peak in SEC corresponding to tetrameric protein and deflavination for AvLCTO using same protocols used for PaLCTO was not successful. Other methods such as urea induced denaturation resulted in protein aggregates upon urea removal, precluding similar studies with AvLCTO. Consistent with other observations, the melting temperature of AvLCTO was higher (Tm 59°C) compared to PaLCTO (Tm 47°C) with a Tm difference of 12°C (S3 Fig). These results implicate a more stable quaternary structure in AvLCTO than PaLCTO and the cofactor dependent assembly of enzyme seems to be a specific property of the PaLCTO enzyme.

### Crystal structure of WT PaLCTO

We solved the crystal structure of PaLCTO with 1.9 Å resolution (Table 1). Crystals used for the study are shown in S4 Fig. The asymmetric unit contains one molecule and the monomer has a typical TIM barrel structure containing eight alpha-helices and beta-strands. Superimposition of WT PaLCTO and AvLCTO monomer shows that the structures are highly similar (rmsd = 0.67 Å). The main structural differences between the monomers of WT PaLCTO and AvLCTO can be observed in the N- and C-termini (Fig 5A). Firstly, there is a small additional β-sheet composed of two strands formed by the residues 2-9 in the N-terminus of PaLCTO. Secondly, the C-terminus of PaLCTO is three residues shorter compared to AvLCTO.

**Fig 5.**
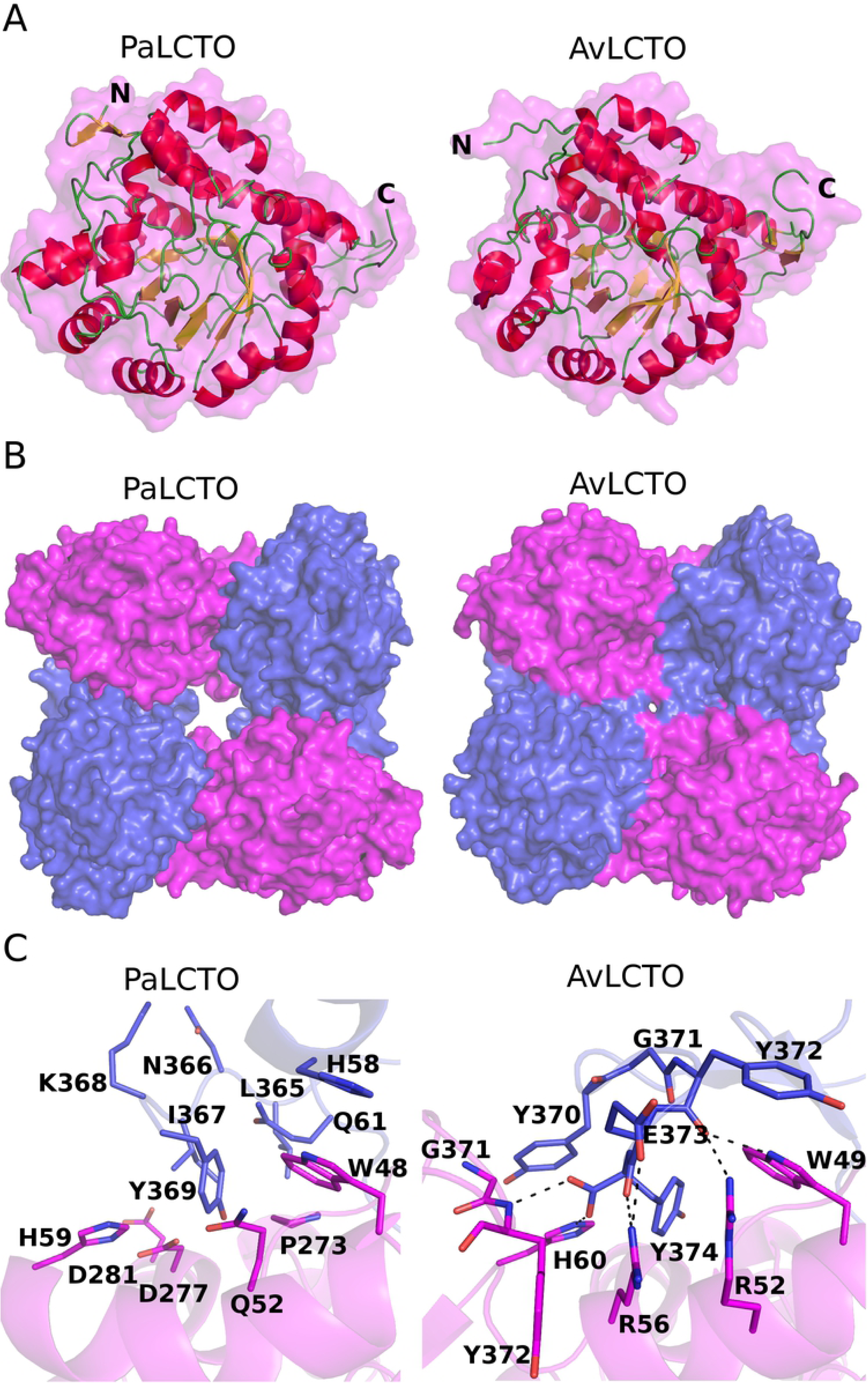
Crystal structure of PaLCTO and structural comparison of PaLCTO and avLCTO tetramers. A) PaLCTO and avLCTO monomers are presented as ribbon-surface models. Beta-strands and alpha-helixes are colored in yellow and red, respectively. Loops and protein surface are colored in green and magenta, respectively. The N- and C-terminus are indicated with the corresponding letters. B) PaLCTO and avLCTO tetramers are presented as surface models. Protein molecules are colored in magenta and blue in turn. C) The interface areas of avLCTO and PaLCTO monomers. The residues are presented as cartoon and stick models and also colored as in B. Hydrogen bond interactions are indicated with black dash lines.

We then analyzed the structure with Pisa server which gave a biological assembly likely similar to AvLCTO. These results are consistent with other data presented above that the protein is tetrameric in solution. Analysis of PaLCTO tetramer with SAXS data measured in solution had a chi^2^ of 1.6 (S5 Fig) which gives additional confirmation that the protein is a tetramer. Structural comparison of the PaLCTO and AvLCTO tetramers shows a similar assembly but it also reveals that the PaLCTO tetramer has a much larger central cavity than AvLCTO due to the shorter C-terminus (Fig 5B). The buried surface area between monomers of AvLCTO is larger (1737 Å^2^) when compared to (1452 Å^2^) from PaLCTO. The C-terminus of AvLCTO contributes to the oligomer formation through a network of ionic interactions (C-terminus, Glu373, Arg52, Arg56), through hydrophobic contact and multiple hydrogen bonds (Fig 5C). The sidechain of Tyr370 stacks with the imidazole ring of His60 from the neighboring subunit and makes the central cavity smaller compared to PaLCTO (Fig 5C). These structural features provide a rationale as to why AvLCTO has higher stability than PaLCTO.

### The active site of WT PaLCTO and the role of the main polypeptide chain S201 – A219

The hydrogen bonding network between FMN and the surrounding residues contains in total twelve interactions (Fig 6A). The 2Fo-Fc electron density map for FMN is shown in S6A Fig. We also observed a glycerol molecule from the cryoprotectant solution bound close to FMN in an expected substrate binding site, where pyruvate (product of the enzyme) has been modelled [21]. Comparison shows that the active site residues of PaLCTO and AvLCTO are highly conserved. All the residues surrounding the co-factor are same and in similar conformation (Figs 6B and 6C) [12,21].

**Fig 6.**
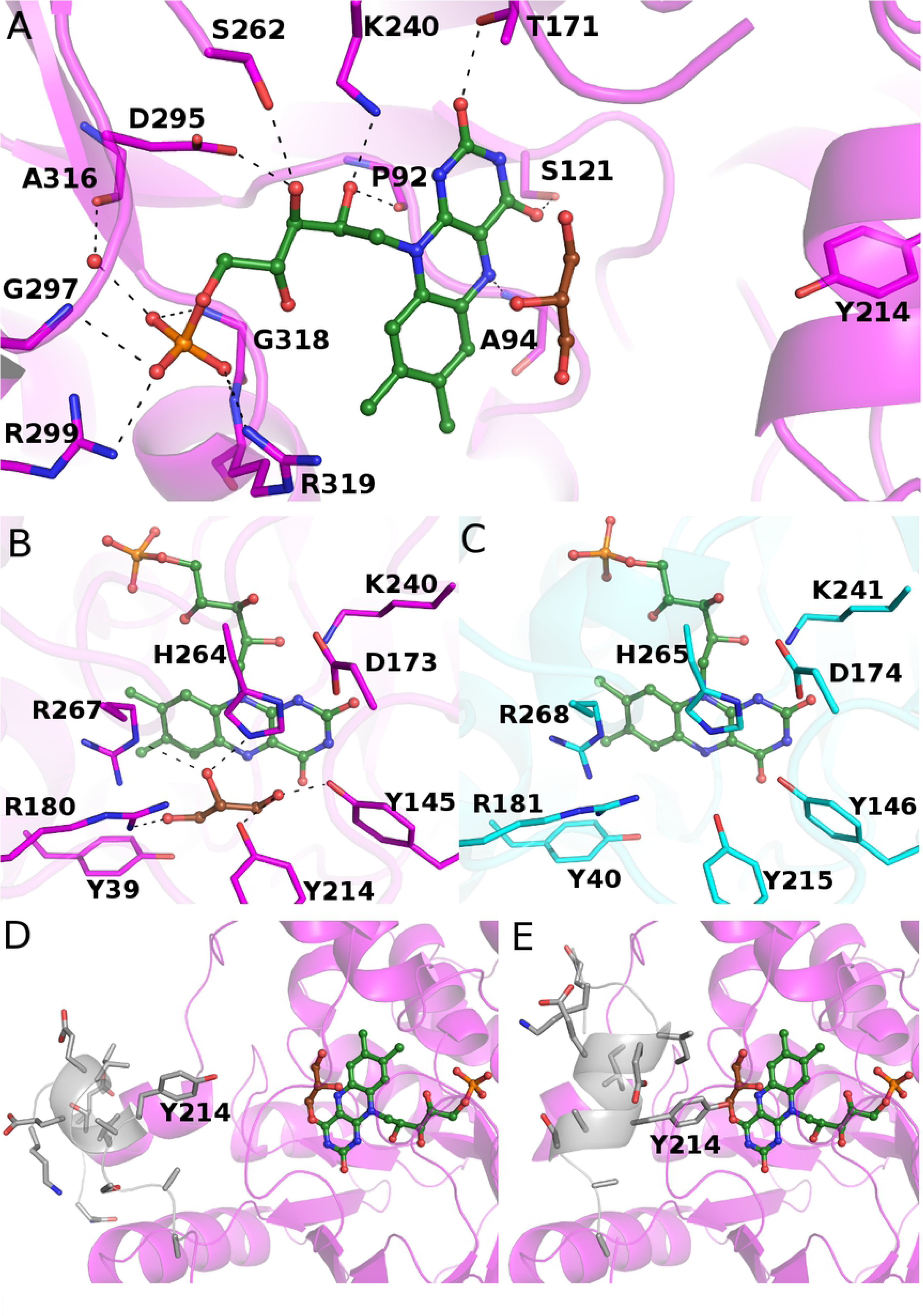
Binding of FMN in WT PaLCTO. A) Hydrogen bonding network between flavin and WT PaLCTO. Flavin and glycerol are presented as stick-sphere models and colored in green and brown, respectively. The residues of WT PaLCTO are presented as cartoon-stick models and colored in magenta. The hydrogen bonds between the flavin and residues are indicated with black dash lines. B) Active site of PaLCTO colored as in A. C) Active site of AvLCTO. The residues are colored in cyan. D) Open form WT PaLCTO. The main polypeptide chain having two conformations is colored in grey and presented as ribbon-stick model. Tyr214 is indicated with a label. E) Closed form of WT PaLCTO. The main polypeptide chain and Tyr214 are presented as in D.

Crystal structure of AvLCTO in complex with the lactate oxidase product pyruvate has been determined earlier [21] and it has been described that the key residues for catalysis are Tyr40, Tyr215, His265 and Arg268. A conserved catalytic triad of (Tyr40, Tyr146, Arg268) positions L-lactate in the active site, Tyr40 and Arg268 interact with the carboxyl group of the substrate, and Tyr146 and Tyr215 interact with the keto group of the lactate C2 carbon. The glycerol observed in the PaLCTO active site mimics the substrate and interacts with these residues (Tyr145, Arg180, Tyr214, His264 and Arg267).

Recent experimental evidence has shown that Tyr215 controls entry and exit of substrate and product in AvLCTO as mutations in Tyr215 result in a change in time of product release [22]. The corresponding residue in PaLCTO is Tyr214 and due to similarity, the loop could have a similar role in this enzyme. Interestingly, we observed that the main polypeptide chain between Ser201 – Ala219 of PaLCTO (loop 4) exists in two conformations (Figs 6D and 6E) in the crystal structure. Because of lack of electron density, we were not able to build both conformations completely and therefore the open conformation lacks the residues Gly205-Gly207 and the closed conformation lacks the residues Ser201-Asn204 in the model. Despite the incomplete structure of the chain, the role of the region could be seen as controlling substrate entry and product release by opening and closing the active site of PaLCTO. In the closed conformation, the OH group of Tyr214 comes very close to the expected substrate having 2.6 Å distance to the O1 atom of the glycerol (Fig 6E). Instead in the open conformation, the distance between the OH group and the O1 atom is 7.5 Å (Fig 6A & Fig 6D).

### Structure of PaLCTO mutant A94G

We generated a mutant (A94G) based on literature precedent suggesting that the mutation of the residue may convert the enzyme to accept also larger substrates [23]. The mutant was crystallized in the identical conditions as WT PaLCTO and the structure was determined at 2.0 Å resolution. The superimposition of the mutant and WT monomer structures showed high structural identity (rmsd = 0.11 Å). However, in the mutant structure we only observed the open conformation for the main polypeptide chain (loop 4: S201 – A219) and we were able to build the conformation completely with all the residues. 2Fo-Fc electron density map of loop 4 (Ser201-Ala 219) is shown in S6B Fig. These results provide an interesting possibility to experimentally determine if A94G mutation reduces loop dynamics for future studies. A stereo view of the A94G mutant aligned to WT PaLCTO is shown in (S7B Fig).

### Structure of refolded PaLCTO

Since refolded PaLCTO behaved similarly in above described experiments, we subjected refolded PaLCTO to crystallization trials and obtained crystals in conditions described for WT native enzyme. The structure was solved to 2.60 Å resolution. A stereo view of refolded PaLCTO aligned to WT native PaLCTO is shown in (S7A Fig). The structure proved that PaLCTO gains the same three-dimensional structure as WT PaLCTO when refolded. We also saw signs of two conformations for the chain S201 – A219 in the structure of refolded PaLCTO, but due to modest resolution and lack of electron density the chains could not be built. Like WT enzyme the refolded oligomer fitted well to the SAXS data (S5B Fig).

### PaLCTO appears not to be a L-lactate oxidase

Activity assays carried out for PaLCTO were surprising as when conducted with L-lactate as substrate, we could not observe any activity although substrate concentrations up to 62.5 mM (∼ 100-fold to wild type K_m_ of AvLCTO) and enzyme concentrations up to 250-fold to those used for AvLCTO. Initially we used His tag cleaved PaLCTO for activity assays and subsequent tests with PaLCTO that was expressed and purified without any tags gave identical results. On the other hand, recombinant AvLCTO produced in-house was active displaying nearly identical activity and K_m_ values compared to commercial AvLCTO (S7 Fig). Km values were 0.519 mM and (in-house produced) and 0.532 mM (commercial enzyme), and V_max_ values were 2926 units/min/mg and 3443 units/min/mg, respectively. These results were puzzling and therefore a commercially available *Pediococcus sp.* lactate oxidase was used as a control and interestingly, commercial *Pediococcus sp* lactate oxidase was active in our assays. To explain this observation, we performed SDS-PAGE of commercial enzyme. Bands were excised and analyzed using mass spectrometry and the commercial PaLCTO was in fact identified as AvLCTO. This was further confirmed by analyzing a separate lot of the commercial enzyme with same results. We used in-house produced AvLCTO and PaLCTO as controls for mass spectrometry to see if it could identify them and they were distinguished correctly. It is likely that a mistake had been made long time ago when the microbial species was identified and has persisted ever since from commercial sources. These results indicate that commercially sold enzyme at least from this specific commercial source is AvLCTO. Additionally, we also tested other substrates such as glycolate and 4-hydroxy mandelate which gave no activity.

Lactate 2-mono-oxygenase is an enzyme with very similar structure and reaction mechanism to lactate oxidase [5]. Therefore, we assayed also lactate 2-mono-oxygenase activity of the in-house produced PaLCTO. These results were negative as there was no acetic acid formed from L-lactate, and neither did it react into any other product, since the concentration of L-lactate was unchanged in the reaction mixture (S9 Fig).

## Discussion

We started with the aim of characterizing recombinant PaLCTO enzyme. Although successful in producing the protein and determining the structure, we were not able to observe enzymatic activity when L-lactate was used as substrate. Given that these enzymes were produced with affinity purification tag, we wondered if this led to loss of enzymatic activity. We also produced recombinant tag free enzyme which gave no activity. As a control we used in-house produced AvLCTO with both affinity tag (tag cleaved later during purification) and tag free version. Both of these versions of AvLCTO had enzymatic activity similar to earlier reports [22]. We also tested other substrates such as glycolate and 4-hydroxy mandelate with PaLCTO but no activity could be seen. Additional experiments for testing lactate monooxygenase activity also failed to detect any acetate as product or disappearance of L-lactate. While this study was in progress a paper describing structure of lactate monooxygenase from *Mycobacterium smegmatis* (MsLMO) was published [24]. Oxidase and monooxygenase enzymes have similar mechanism (see introduction), but monooxygenase enzymes undergo ‘coupled’ pathway. In *M. smegmatis* and other lactate monooxygenases loop 4 is significantly longer and forms a defined folded structure in MsLMO (PDB:6DVI). In MsLMO crystal structures, loop 4 is about 49 residues long and structured elements in the loop have been hypothesized to impede loop 4 dynamics, thereby oxidizing pyruvate further to acetate. Unlike other oxidases including PaLCTO structure, loop 4 in MsLMO is clearly visible, indicating that the loop is less dynamic than in other oxidases. Clearly loop 4 in PaLCTO is more similar to AvLCTO than MsLMO. Given that the active site residues are conserved, and the loop 4 structure is similar to AvLCTO, PaLCTO would catalyze a similar reaction as AvLCTO, although the identity of the substrate is not known. Commercially available PaLCTO turned out to be AvLCTO and further studies are required to definitively specify the origin of the enzyme. These results could indicate that *P. acidilactici* does not have any active lactate oxidase but does not rule out from other species.

We report to our knowledge the first protein that can reversibly form oligomers in presence of FMN, while several other examples of FAD containing proteins exhibit this behavior [8,19,20]. This observation was specific to PaLCTO as it was not observed for AvLCTO. Based on structural studies we believe that the C-terminal residues provide inter subunit interactions that stabilize AvLCTO tetramer and the absence of such interactions makes PaLCTO susceptible to changes in oligomeric state. An interesting observation with deflavinated monomeric PaLCTO was that although it lacked any secondary structure elements it was still globular based on SAXS data. One useful feature of PaLCTO, is that there are no cysteines in the wild-type protein. This allows one to introduce site-specific cysteines that could allow cysteine labelling by fluorophores and other small molecules to monitor protein folding. With high expression levels, protocols for folding and unfolding and availability of crystal structure, makes PaLCTO potentially an attractive model protein to study protein folding.

## Accession codes

Atomic coordinates and structure factors have been deposited to Protein Data Bank with accession codes 6RHT (WT), 6RHS (refolded) and 6RHV (A94G mutant).

## Acknowledgements

The use of the facilities of the Biocenter Oulu for DNA sequencing, Proteomics and Protein Analysis and Protein Crystallography core facility, a member of Biocenter Finland and Instruct-FI, are gratefully acknowledged.

## Funding Sources

This work was funded by Academy of Finland (grant no. 287063 and 294085 for LL) and by European Regional Development Fund (LIIKUTPA project, Kainuun Liitto, TK and PK). The funders had no role in study design, data collection and analysis, decisions to publish, or preparation of the manuscript.

## Supporting information

**S1 Fig. Purification of PaLCTO.** A) Chromatogram obtained during preparative size exclusion chromatography. T and M indicate tetrameric and monomeric fractions. Peak labelled T was yellow in color and Fraction M was colorless.

**S2 Fig. SAXS Guinier plots.** A) Guinier fit for native WT PaLCTO B) Guinier fit for monomeric PaLCTO C) Guinier fit for refolded PaLCTO.

**S3 Fig. CD studies for PaLCTO and AvLCTO.** A) Comparison of native and refolded PaLCTO. B) CD spectra of refolded PaLCTO at 22 and 70°C. C) CD spectra of avLCTO at 22 and 86°C. D) Melting curve analysis of avLCTO using parameters obtained from 222 nm. Melting temperature was 59°C.

**S4 Fig. Crystals of PaLCTO.** Thin square plate-like crystals were used for X-ray diffraction studies.

**S5 Fig. CRYSOL fitting of experimental protein structures into SAXS curves.** A) Analysis of PaLCTO WT structure with raw scattering curve B) Analysis of refolded PaLCTO structure with experimental SAXS data.

**S6 Fig. 2Fo-Fc map of relevant regions.** 2Fo-Fc electron density is shown in blue and contoured at 1 σ. a) FMN density in WT native PaLCTO b) Loop 4 (Ser201-Ala219) electron density in A94G mutant is unambiguous and clearly visible. Gatekeeper residue Tyr 214 is also shown.

**S7 Fig. Stereo view of structures.** A) Alignment of WT native PaLCTO and refolded PaLCTO B) Alignment of WT PaLCTO and A94G mutant PaLCTO. Loop 4 in WT and A94G mutant is shown in Sticks. In both the panels WT PaLCTO is shown in orange.

**S8 Fig. Comparison of activity produced in-house and commercially available AvLCTO.** A) Michaelis-Menten kinetics of commercially available AvLCTO. B) Michaelis-Menten kinetics of in-house produced AvLCTO. V_max_ and K_m_ values are shown in respective graphs.

**S9 Fig. L-lactate appears not to be a substrate of annotated *Pediococcus* lactate oxidase**. In-house produced PaLCTO and commercial AvLCTO were incubated overnight at +37°C with L-lactate in dimethylglutarate buffer, pH 6.5 using imidazole-HCl, pH 7.0, as enzyme diluent. The reaction mixtures were analyzed by capillary electrophoresis together with a buffer control and standards lactate, pyruvate (product of lactate oxidase reaction) and acetate (product of lactate 2-monooxygenase reaction). Similar experiment was done also using Na-phosphate or Hepes buffer as enzyme diluent with equal results. For clarity, the buffer control and lactate standard graphs are not presented in the figure. Inserted table shows lactate concentrations after overnight incubation. The capillary electrophoresis graphs indicate catabolism of L-lactate by AvLCTO, and inactivity of PaLCTO towards L-lactate at least in these conditions. The formed pyruvate is under the dimethyl glutarate peak, whereas acetate, if any were generated from enzymatic reactions, should be visible. In-house produced AvLCTO was used in some experiments and displayed equal activity to commercial AvLCTO.

**S1 Table. SAXS summary.**

